# Metagenomic estimation of dietary intake from human stool

**DOI:** 10.1101/2024.02.02.578701

**Authors:** Christian Diener, Sean M. Gibbons

**Author notes:** Correspondence can be addressed to and.

## Abstract

Dietary intake is tightly coupled to gut microbiota composition, human metabolism, and to the incidence of virtually all major chronic diseases. Dietary and nutrient intake are usually quantified using dietary questionnaires, which tend to focus on broad food categories, suffer from self-reporting biases, and require strong compliance from study participants. Here, we present MEDI (Metagenomic Estimation of Dietary Intake): a method for quantifying dietary intake using food-derived DNA in stool metagenomes. We show that food items can be accurately detected in metagenomic shotgun sequencing data, even when present at low abundances (>10 reads). Furthermore, we show how dietary intake, in terms of DNA abundance from specific organisms, can be converted into a detailed metabolic representation of nutrient intake. MEDI could identify the onset of solid food consumption in infants and it accurately predicted food questionnaire responses in an adult population. Additionally, we were able to identify specific dietary features associated with metabolic syndrome in a large clinical cohort, providing a proof-of-concept for detailed quantification of individual-specific dietary patterns without the need for questionnaires.

## Introduction

Dietary intake and nutrition are key determinants of human growth and development, metabolic health, and chronic disease risk ^1–3^. Diet also shapes the composition of the human gut microbiota ^4^, and in turn, the effects of diet on the host can be influenced by the ecology of the gut ^5^. Across the lifespan, dietary intake patterns can either alleviate or exacerbate a wide range of disease conditions, including cardiovascular disease, diabetes, Alzheimer’s disease, and cancer ^6–9^.

Accurate tracking of dietary intake, including the quantification of dietary metabolites and energy content, is critical to understanding phenotypic heterogeneity in human cohort studies. However, diet tracking is often hampered by challenges in obtaining high quality, unbiased data on dietary intake ^10^. In many cross-sectional studies, individual dietary intake data are obtained from questionnaires, which need to strike a balance between granularity and ease-of-use. The most common methodology employed for assessing dietary intake, the food frequency questionnaire (FFQ), asks participants for coarse-grain information on habitual dietary patterns for a set of common food categories ^11^. More detailed information can be obtained from dietary recall surveys, where study participants fill out food diaries, sometimes with clinician-support, for a pre-specified time frame ^12^. Dietary recall data provides more detailed quantification of food intake and can be used to generate more detailed mappings to macro- and micronutrient intake, but these methods are often dependent on proprietary platforms and strong participant compliance, making it difficult to compare dietary intake data across cohorts and to maintain participant compliance in longitudinal studies ^13^. Standardized questionnaires, or common food databases, often do not reflect diverse human populations, leading to inequalities where the diets of minority populations are not accurately represented in existing questionnaires or recall surveys ^14^. Finally, dietary questionnaires rely on the ability of study participants to accurately and without bias recall their own food intake, which is known to be fraught and has led to some debate on the utility of self-reported dietary data ^15–17^. Thus, there is demand for approaches that can quantify dietary and nutrient intake patterns without the need for FFQs or recall surveys.

One survey-free technique for coarsely assessing diet quality is the quantification of major diet-influenced analytes in human plasma or serum. This approach is used widely in clinical settings, where regular measurement of blood glucose, cholesterol, and other lipid levels are the current standard of care ^18,19^. However, these clinical chemistries represent a limited breadth of diet-relevant features that can be accurately quantified, which could be expanded on by using targeted or untargeted metabolomics of blood, saliva, or fecal samples. Another approach has been to capture images of meals (e.g., with a smartphone) and apply machine learning to these images to track dietary intake ^20,21^. Image tracking and physical sensors have proven to be challenging approaches, requiring large training databases, showing a limited ability to estimate portion size, and relying on a fairly high degree of participant compliance ^22,23^.

Molecular ‘omics approaches to diet tracking have the potential to increase the sensitivity and resolution of intake assessments, while reducing the compliance burden for participants. Due to the complexities of absorption and metabolism of the dominant compounds found in the diet, blood metabolomics currently provide limited information on food intake. To overcome this limitation, recent work has focused on constructing curated databases of food-specific MS spectra that can be used to identify the presence of specific foods in fecal or blood samples ^24,25^. These reference spectra can identify specific food components with high accuracy, but they provide limited information about the overall abundance of a food item. An alternative approach is to quantify dietary intake using residual food-derived DNA in stool. Prior studies have shown that plant intake patterns can be quantified in human stool by targeting plant-specific marker genes for amplicon sequencing ^26,27^. While effective, these targeted methods require additional sample processing prior to sequencing. Metagenomic shotgun sequencing (MGS) of stool DNA, on the other hand, is a widely generated data type in human microbiome research ^28,29^. Leveraging MGS data directly for diet tracking would require no additional sample processing steps. However, there are several challenges to detecting food related DNA in MGS data, which has delayed the implementation of metagenomic-based diet tracking.

Fecal MGS data is commonly used to quantify the taxonomic and functional composition of bacterial, archaeal, viral, and fungal communities in the gut, where genes can generally be identified with *de novo* methods ^30,31^. However, while we expect food-derived DNA to be present in stool, the low frequency of these reads relative to microbial- and host-derived reads makes *de novo* gene prediction from these sequences intractable ^32^. Alternatively, individual reads can be mapped to large databases of reference sequences, using efficient hashing schemes, in order to generate read-specific taxonomic annotations ^33–35^. However, quantification of dietary intake via reference-based approaches is hampered by the increased genomic complexity of higher eukaryotes and by the lack of dedicated food genome databases. Furthermore, these reference mapping approaches are prone to false-positive assignments, requiring the inclusion of decoy genomes in the database from other organisms that are known to be present in the sample, like host-associated reads and reads coming from the microbiota ^36,37^. Finally, even if we could quantify the taxonomic composition of food-associated reads, we currently lack the ability to automatically translate this taxonomic information into nutrient content.

Here, we aimed to overcome these limitations by building a comprehensive food genome database for annotating food-associated reads in human stool, along with high-resolution mappings between food items and dietary metabolite profiles. We paired the constructed database with a scalable and decoy-aware mapping strategy and validated its performance using simulated and *in vivo* data from infants and adults, showing that we can accurately quantify dietary components and nutrient intake from MGS data. Finally, we find that our MGS-based dietary assessments were strongly associated with variation in metabolic health in a large European cohort.

## Results

### Linking food genomes to nutrient information

While databases that map food items to nutrient content exist, such as the USDA FoodData Central (https://fdc.nal.usda.gov/) and FOODB (www.foodb.ca), none of these databases are directly linked to genomic data from plants, animals and fungi present in the human diet. 619 of the 992 foods present in FOODB can be mapped to the NCBI taxonomy, and we aimed to obtain genomic assemblies for as many of these mappings as possible. Food items were mapped through NCBI taxonomy IDs in a multi-tiered approach (Fig. 1A). Food items were first matched to RefSeq genomes at the species level and then at the genus level if no match could be found for the respective species, yielding a set of 377 foods mapping to 307 unique genome assemblies (Fig. 1B-C). This was followed by a search in the full NCBI Nucleotide Database on the species and genus level in order to obtain partial assemblies for food items without a full reference assembly. 95 partial assemblies representing 99 additional foods could be identified in this way (Fig. 1B-C), resulting in a final database containing 402 genomes and genomic assemblies representing 476 unique food items (77% of all foods in the FOODB with a taxonomic identifier).

**Figure 1.**
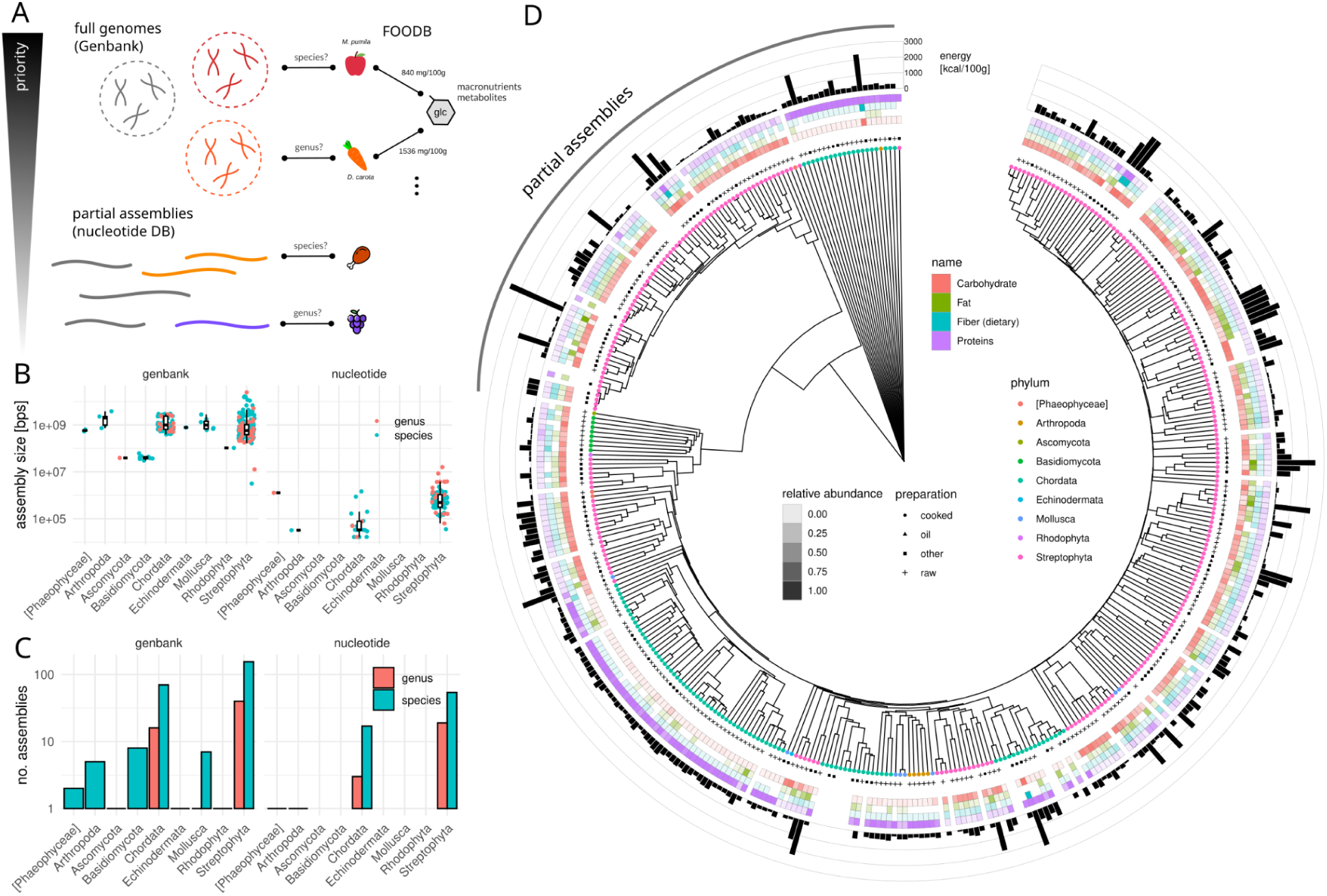
Constructing a metagenomic food database. (A) Illustration of the search strategy used to map food items to assemblies and their connection to nutrient content. (B) Assembly size for the identified food-related organisms. Titles denote the database yielding the hit (Genbank - complete genomes, nucleotide - partial assemblies). (C) Number of food organisms matched and the respective taxonomic rank where the match was found. (D) Phylogenetic tree of the identified food organism assemblies, generated using UPGMA on estimated average nucleotide identity (ANI). Colored circles denote the phylum, symbols indicate the dominant food preparation type, filled rectangles show macronutrient composition per 100 grams of biomass, and black bars show the energy content of individual food-assembly pairings per 100 grams of biomass.

Macronutrient, energy, and specific metabolite contents were collapsed for each matched taxon to the mean composition across the 402 genomic assemblies, while keeping track of the predominant preparation type (cooked, raw, oil, etc). The resulting database contained a total of 476 billion base pairs covering all major phyla of common food components, along with their average nutrient composition (Fig. 1D). The majority of genomic data came from the phylum *Streptophyta*, which includes most of the common plant-based foods, followed by the phylum *Chordata*, which comprises most of animal-derived foods (Fig. 1B,D). Phylogenetic distance, quantified by average nucleotide identity, was associated with relative protein and carbohydrate content of the food items (Fig. 1D, PERMANOVA p=0.001, R^2^=0.06 and 0.05, respectively), revealing that there is a weak phylogenetic association with macronutrient content. In particular, macronutrient composition varied only by 10-20g/100g making within 90-95% ANI identity, suggesting that nutrient composition is similar even when mapping reads only to the closest genus (SI Fig. S1).

### Decoy-aware, efficient mapping allows for accurate quantification of dietary intake

The size of our food genome database exceeded commonly used databases for the classification of bacteria, archaea and viral genomes by at least 5-fold. Thus, we developed a computationally efficient mapping strategy, which we termed Metagenomic Estimation of Dietary Intake (MEDI). MEDI is based on the KRAKEN2 mapping scheme, to ensure scalability to very large data sets ^33^. KRAKEN2 uses a fast kmer hash to identify the least common ancestor (LCA) for each single read, which can be paired with a Bayesian redistribution approach that passes down the phylogenetic tree (BRACKEN) ^38^. Because the majority of genetic material in human-associated stool microbiome samples is likely from bacteria or the host, there is a high chance of false positive identification of background DNA as food components. To avoid this, we opted for a decoy-aware approach, where the kmer hash also included additional background genomes belonging to bacteria, archaea, viruses, common plasmids, and the human genome ^39^. We combined this with an additional post-classification filtering step (prior to BRACKEN redistribution) that removed individual reads with inconsistent kmer classification patterns that included taxa from distant clades (see Fig. 2A and Methods). The resulting abundance estimates were then used to identify the food items present in a sample and to derive the macronutrient and metabolic composition (standardized to a 100g portion of a mixture of the respective food items), where the relative abundance of a given food item in a sample was used to calculate its relative contribution to the dietary nutrient content (see Methods).

**Figure 2.**
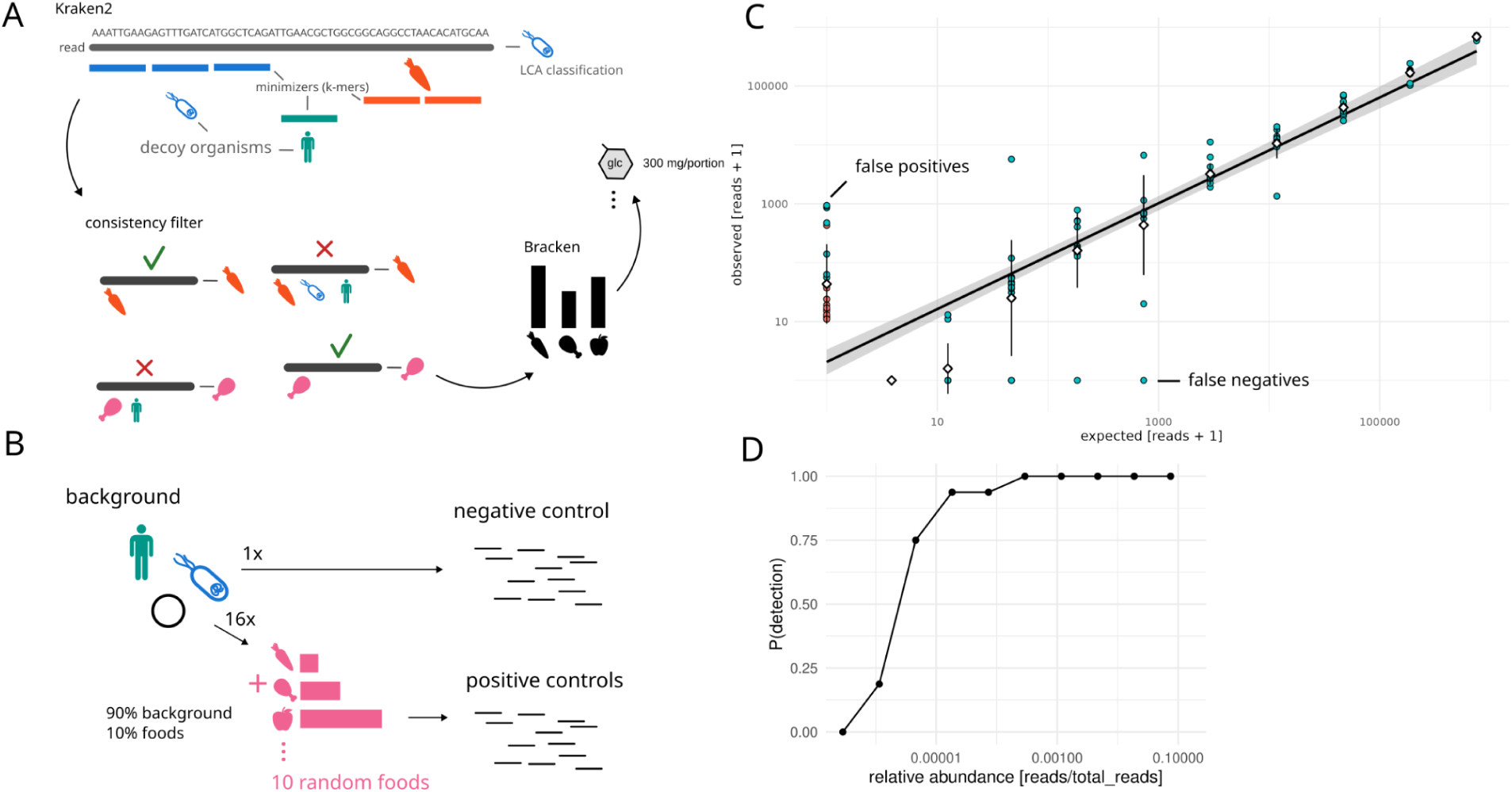
Food genome quantification on simulated ground truth data. (A) Illustration of the mapping and filtering strategy used by MEDI. Individual minimizer assignments (k-mer classifications) were used to assign consistency scores to reads and to filter reads with discordant mappings. (B) Sampling strategy for the ground-truth data. All samples contain at least 90% background of an average bacteria, archaea, and host background. Positive samples contain simulated reads from 10 random food assemblies with logarithmically-varying abundances. (C) Quantification performance across simulated negative and positive controls. Points denote detected food items. Black line denotes a linear regression fit and gray area the 95% confidence interval. Fill color denotes whether the identified food items were phylogenetically close to a ground truth organism (blue, within the same genus) or not (red, no genus overlap). False positive organisms are generally connected to organisms in the same genus. (D) Probability of detecting a true positive food item in a sample as a function of relative food item abundance (i.e., detection power).

MEDI was tested on an artificial ground truth data set using simulated reads. Briefly, we generated an average abundance profile of the decoy organisms present in fecal samples from a healthy population of 351 individuals from the integrative human microbiome project, drawing subsamples from this background distribution to simulate different decoy communities (iHMP) ^28^. Positive control samples (i.e., samples containing a dietary signal) were generated by injecting 10% food reads from 10 randomly chosen food items into each individual sample (Fig 2B). Given the high prevalence (90% of total abundance) and richness of decoy organisms in the simulated data set, one would expect methods that incorrectly map background taxa to foods to perform poorly. The ten reference foods added to each sample were logarithmically staggered in abundance within each sample, creating a relative abundance range of 0.00003-7.5% across food items. A background sample without any food reads added was used as a negative control. This simulated data set allowed us to assess the prevalence of true positives, false positives, true negatives, and false negatives, as well as to quantify the taxonomic specificity and detection limits of our approach. MEDI was able to accurately identify and quantify dietary intake in all simulated samples (Fig. 2C, mean R^2^=0.9, p<10e-6 for all positive samples). Only 13 of the 10 million reads in the food-negative sample were classified as food-derived. The false positive rate was slightly higher in the food-positive samples, where misidentified food items were generally from the same genus as a more abundant true positive food item also present in a sample, which did not alter quantification accuracy for the true positive food items (Fig. 2C). MEDI was highly sensitive, providing >80% power for detecting a food item with a relative abundance as low as 0.001% (10 reads per million, see Fig. 2D). In summary, MEDI was able to accurately distinguish and quantify food items in simulated metagenomic samples, with a negligible rate of cross-domain mismatching from the gut microbiome or the host.

### Applying MEDI to infant and adult stool metagenomes with paired dietary questionnaire data

To assess the frequency of food-derived reads across different stages of life, we quantified food abundances and contents in fecal samples using MEDI across two large human data sets from infants and adults. Infant metagenomic shotgun sequencing data were obtained from a previously published cohort from St. Louis, USA, describing 447 longitudinal fecal samples from 60 infants of 1-253 days of age ^40^. Adult reference samples were obtained from 351 healthy individuals (mean age of 31 years) within the iHMP project, as this subcohort includes standardized food frequency questionnaires ^41^.

As expected, we generally observed a lower prevalence of food-derived reads in infant stool than in adult stool. Food-derived genomic material could be detected in less than 50% of infant fecal samples up to the onset of solid food intake (around day 160), where the prevalence of samples with detected food reads increased steadily (see Fig. 3A). This presence-absence pattern was not correlated with age or feeding type (breast-fed or formula, logistic regression p>0.1). In contrast, food-derived genomic material was detected in 99% of adult samples (361/365). Total relative abundances of detected food items varied across infant samples, and increased with age independent of bacterial biomass and delivery mode (repeated measures mixed effects model, log beta=0.11, p=0.0003), with a notable increase at the onset of solid food consumption (Fig. 2B). A somewhat higher relative abundance of foods could be observed in the first 3 days of life, but this signal was accompanied by a smaller fraction of bacterial reads (especially in infants delivered via Cesarean; see red points in Fig. 2B), which increases the sensitivity for food detection. Bacterial biomass remained stable after the first 3 days of life in infants and in adults (usually representing more than 99% of metagenomic reads, Fig. S2). In contrast to total bacterial relative abundance, which differed very little across samples (Fig. S2), total food-read relative abundance varied over three orders of magnitude in both infants and adults (0.007-1.3% of total reads). In summary, while the relative metagenomic abundances of bacterial or human reads are generally quite stable, food-read relative abundances are much more variable across infants and adults.

**Figure 3.**
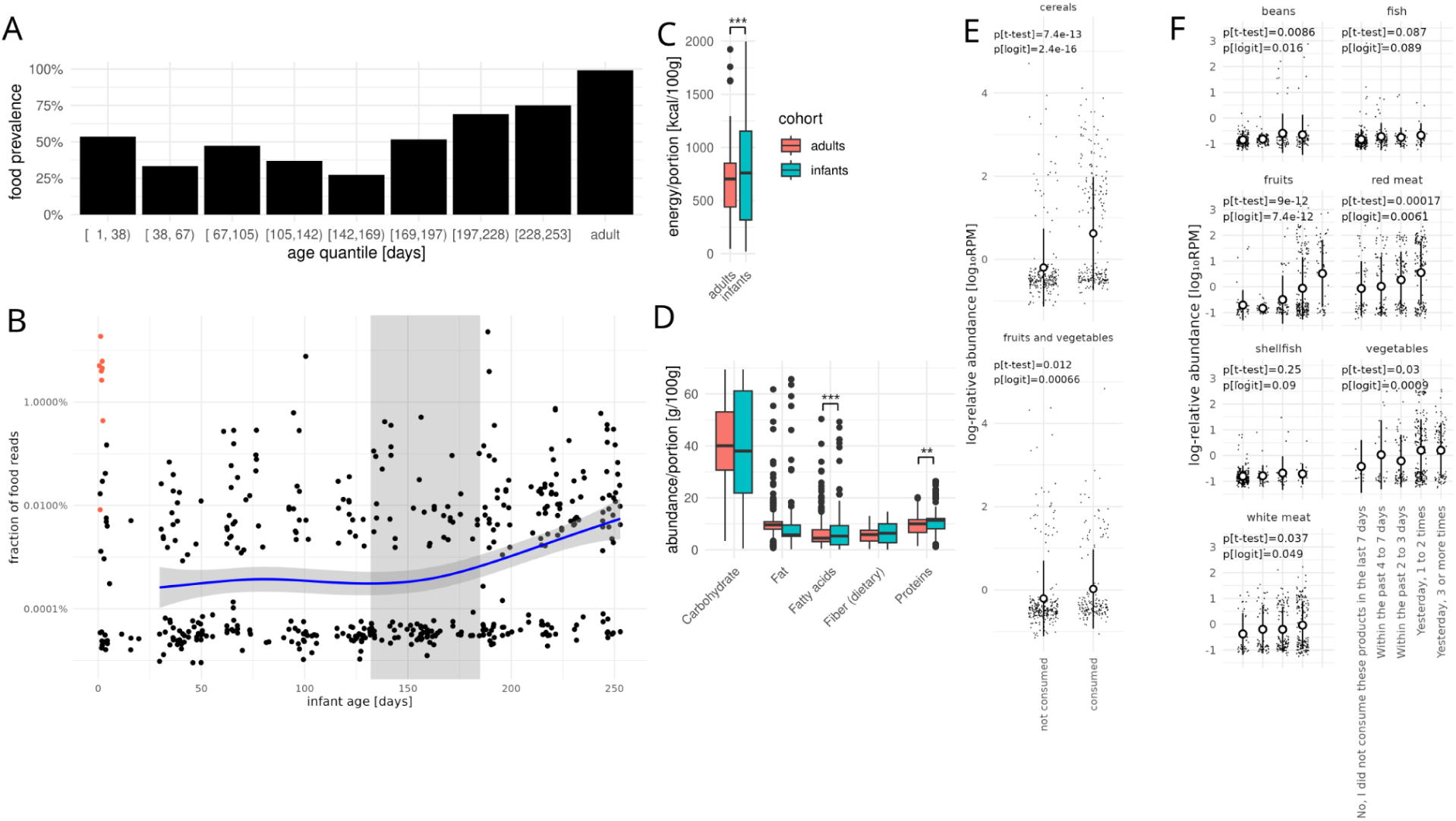
MEDI food abundances across infants and adults. (A) Fraction of samples with at least one detected food read across different age groups. (B) Relative abundance of food-derived reads in a cohort of 447 infants. Blue line denotes the smoothing spline of the observed reads. Light blue area denotes the 95% confidence interval of the spline curve. Orange dots denote samples with less than 95% overall abundance mapped to bacteria (i.e., low bacterial biomass). Gray shaded area denotes the interquartile area of the onset of solid food intake across infants. (C) Energy content per standardized portion size (100g) per sample in adults and infants. Stars denote significance: * - p<0.05, ** - p<0.01, *** - p<0.001 (D) Macronutrient content per standardized portion size in infants and adults. Stars denote significance: * - p<0.05, ** - p<0.01, *** - p<0.001. (E) Comparison of relative food group abundances with paired diet frequency questionnaire data from infants. (F) Comparison of MEDI-predicted relative food group abundances with diet frequency questionnaires in adults. In (E-F) p[t-test] indicates the p-value of a Welch t-test of log-transformed relative abundances and p[logit] denotes the p-value of a logistic regression of food occurrence against food frequency strata.

Mapping the nutrient and metabolite composition of the identified foods to a standardized portion allowed us to compare nutrient composition between infants and adults. Energy content per portion (kcal/100g) was positively correlated with infant age (r=0.33, Pearson test p<10e-6) in concordance with an increased calorie demand for growth across the infant lifespan and with an increased reliance on solid foods with infant age. Similarly, the average energy density of the MEDI-inferred diets was higher in infants than in adults (Fig. 3C, Welch t-test p=0.00003). Macronutrient composition was similar between infants and adults and mirrored common nutritional recommendations ^42^. Compared to infants, adult diets contained less protein (Welch t-test p=0.0011) and fewer fatty acids per standardized portion (Welch t-test p=0.0008).

MEDI-inferred dietary intake was concordant with food frequency questionnaire data from both infants and adults. Reported consumption of fruits, vegetables, and cereals led to increased prevalence (logistic regression) and abundance (linear regression) of food-derived reads in infants (see Fig. 3D). Within the adult iHMP cohort, food frequency patterns were captured by MEDI estimates of prevalence and abundance for several food categories that could be mapped to FOODB food groups or subgroups (Fig. 3E), with the exception of fish and shellfish, which generally showed relative abundances beneath the MEDI detection limit (i.e., <10 reads per sample, on average; Fig. 2D).

In summary, MEDI was able to accurately capture dietary and nutritional intake across infants and adults, directly from stool MGS data.

### Applying MEDI to a cross-sectional study of dietary patterns in patients with and without metabolic syndrome

To illustrate the ability of MEDI to identify dietary patterns associated with health and disease states, we performed a prospective cross-sectional study to identify dietary features associated with metabolic syndrome in the absence of dietary questionnaire data. Here, we leveraged a subcohort of 533 individuals with paired fecal samples and metabolic health information from the METACARDIS study ^43^. The selected cohort consisted of 274 healthy individuals (HC in METACARDIS), and 259 individuals with varying clinical manifestations of metabolic syndrome, split into 134 individuals receiving medication (MMC in METACARDIS) and 125 untreated individuals (UMMC in METACARDIS). MEDI identified a mean of 4805 food-derived reads per sample (141-589,755, Fig. 4A). Wheat, hibiscus, cocoa, pork, oats, and flax were the most commonly detected food items, accompanied by many food items that were only detected in a small subset of the cohort (Fig. 4A). Similarly, MEDI-inferred macronutrient and metabolite composition varied substantially across individuals, with a set of highly prevalent compounds detected across the cohort and other sets of metabolites only observed in small subgroups of individuals (Fig. S3). Nutritional profiles could be clustered into a smaller subspace of carbohydrate and protein content, with a tendency towards higher energy content in high protein/low carbohydrate diets (Fig. 4B). However, neither protein nor carbohydrate content were associated with metabolic syndrome (Welch t-test of HC vs. MMC/UMMC, both p>0.05).

**Figure 4.**
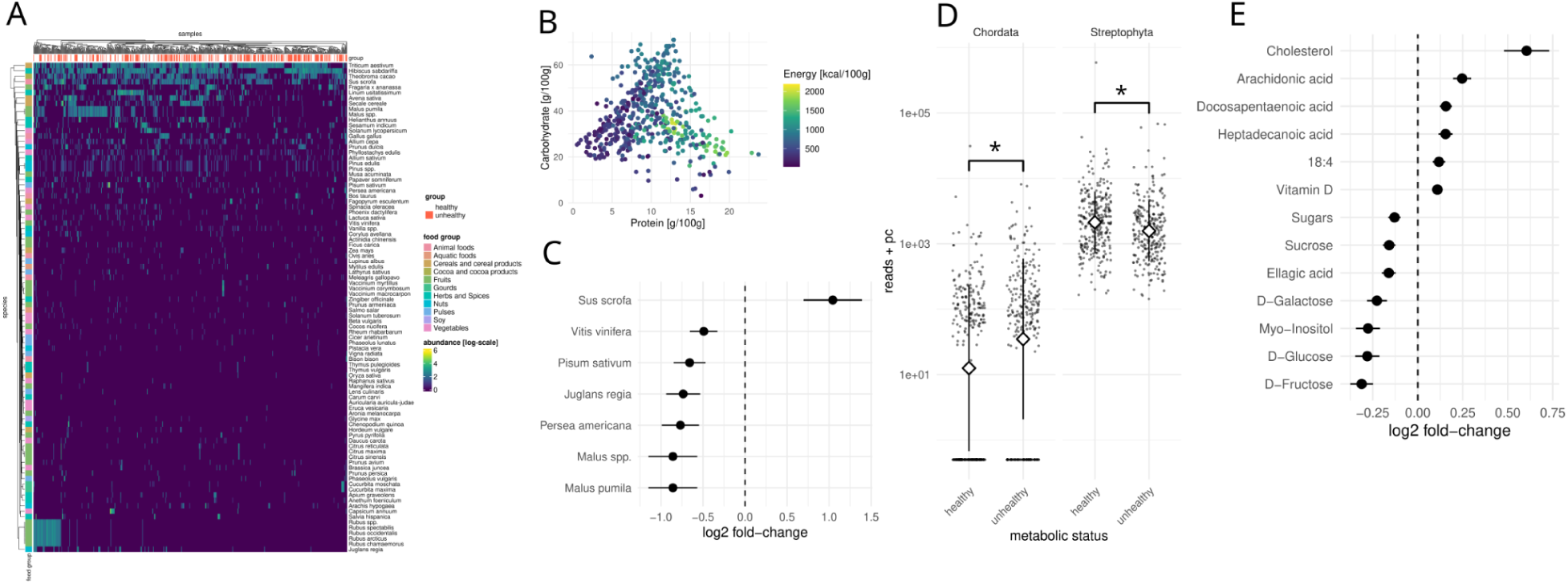
MEDI dietary intake estimates were associated with metabolic health. (A) MEDI-detected food abundances across a cohort of 533 metabolically healthy and unhealthy individuals from the METACARDIS cohort. Fill colors denote abundance (log10[reads + 1]). Column annotations denote metabolic health status from the original METACARDIS cohort. Row annotations denote the major food groups from FOODB. (B) Relationship between protein and carbohydrate abundances for all samples. Fill color denotes energy content. (C) Food-derived organisms with a significant association with metabolic health (FDR-corrected p<0.05 in a LIMMA-VOOM regression of read counts vs. metabolic health status). Bars denote standard errors. (D) Food-derived phyla associated with metabolic health. Stars denote FDR-corrected LIMMA-VOOM p-values (* - p<0.05). (E) Food-derived compounds associated with metabolic health (FDR-corrected p<0.05 in a linear regression of log abundance vs. metabolic health status). Bars denote standard errors. In (C, E) positive log fold-changes denote increased abundances in metabolically unhealthy individuals and negative log fold-changes denote species more abundant in healthy individuals.

We next ran a systematic differential abundance analysis to identify dietary patterns associated with metabolic syndrome (MMC/UMMC) states, compared to healthy individuals (HC). We found that fecal samples from individuals with metabolic syndrome contained approximately twice as much pork as samples from healthy individuals, which in turn contained more fruit (apple, grape, pineapple, avocado), pea, and walnut DNA (Fig. 4C, LIMMA-VOOM log-normal regressions, all FDR-corrected p<0.05). On a broader scale, metabolic syndrome was also associated with a higher abundance of *Chordata* reads, denoting vertebrate animal-derived foods, and a lower abundance of *Streptophyta*, which includes most plants found in the human diet. These results are consistent with prior studies that have identified higher consumption of animal products, such as pork, and a lower consumption of fruits and vegetables, as risk factors for metabolic syndrome and cardiovascular disease ^44,45^.

Several macronutrient and metabolite abundances (per standardized portion) were associated with metabolic syndrome or health in this cohort. Cholesterol abundance per portion was around 4 times higher in individuals with metabolic syndrome, as were several omega fatty acids and vitamin D. Diets from healthy individuals contained higher abundances of sugars, myo-inositol, and ellagic acid. Though it seems counterintuitive that higher sugar consumption protects against metabolic syndrome, one should note that MEDI quantifies only non-supplemented (i.e., non-refined) sugars which, for the most part, are coming from fruits and other plants. Consequently, MEDI was able to identify dietary patterns known to be associated with metabolic syndrome in a questionnaire-free setting, directly from stool MGS data.

## Discussion

Obtaining accurate and unbiased estimates of dietary intake from large cross-sectional and longitudinal human cohorts is a fundamental challenge that has yet to be resolved. Here we introduced MEDI, a method for quantifying dietary intake directly from human stool DNA. Metagenomic shotgun sequencing of stool DNA is the current gold standard in quantifying the taxonomic and functional composition of the human gut microbiome and MEDI makes it possible to derive dietary intake data from this widely-available data type. MEDI will allow for the extraction of dietary information from the hundreds of thousands of human stool metagenomic samples that have been deposited in public databases. Dietary patterns are a major determinant of microbiome composition and a strong confounder of human cohort studies. Thus, we feel that MEDI will be a valuable tool for nutritionists, epidemiologists, anthropologists, clinicians, and microbiome researchers.

The total relative abundances of food-derived reads varied greatly across infants and adults, ranging from 0 to 15% of the total sequencing reads. This stands in stark contrast to bacterial or host-derived (human) reads, whose relative abundances do not vary much across individuals (Fig. S2). This is likely a consequence of the dynamic nature of food items passing through the human body, where DNA from whole foods, especially from plants, is more likely to survive passage through the gut than DNA from ultra-processed foods. For example, the low prevalence of food DNA in infant samples could be due to breast milk intake (i.e., we do not consider human DNA as a component of the diet) and the consumption of infant formula (i.e., highly-processed food). Despite this inherent variation, the relative abundance of food-derived DNA was strongly associated with age in infants and mirrored the transition to solid foods. Food DNA was found in most adult stool metagenomes (>99%) and quantitatively recapitulated food frequency questionnaire data (Fig. 3). However, while food frequency questionnaires only allowed for the quantification of broad food groups, MEDI could identify individual species and distinguish between differences in food prevalence (consumption frequency) or abundance (relative amount consumed).

Additionally, MEDI allows for the mapping of identified food items to nutrient and metabolite intake, given a representative portion. These estimates of nutritional intake stood in good agreement with general nutrition recommendations and were able to identify anticipated shifts in nutrient consumption between infants and adults ^46,47^. Finally, MEDI-inferred nutritional intake was consistent with dietary markers of metabolic syndrome, even in the absence of questionnaires or metadata (Fig. 4).

Despite the promise of DNA-based dietary tracking there are certain limitations to these approaches that merit discussion. For example, many commonly consumed food items, including highly processed foods or supplements, do not leave a residual DNA signal in stool, which biases DNA-based detection towards unprocessed whole foods (although see discussion above about putative detection of DNA from common food additives). This bias extends to the subsequent nutrient mappings, which do not account for common dietary ingredients, like processed sugars or cooking oils. This is reflected in the observed association between MEDI-derived sugar abundances and better metabolic health, due to the fact that MEDI-predicted sugars are largely derived from fruit ^48,49^. However, there may be merit in disentangling sugars from whole foods, such as fruits and vegetables, from processed sugar. Most processed sugar is likely absorbed in the small intestine. Metabolites associated with fecal DNA may have survived passage through the upper gut, and are more representative of the nutrient environment in the colon, which is particularly relevant to the metabolic activity of our commensal microbiota. MEDI may also provide insight into the consumption of certain highly-processed foods, due to the presence of DNA that is likely derived from common non-caloric bulking agents added to these foods, like cotton-derived cellulose and pine wood pulp ^50,51^. Specifically, hibiscus (cotton plants are closely related to hibiscus) and pine tree DNA were among the most prevalent food components identified in MEDI-inferred diets from the METACARDIS cohort. While hibiscus flowers and pine nuts are also present in the diet, cross mapping of reads from cotton and pine wood pulp can occur. Another limitation is that existing food databases are biased towards the diets of largely white, affluent populations in Europe and America, often lacking foods consumed in indigenous societies, in non-industrialized countries, or within minority populations in industrialized countries ^52,53^. This is part of a larger issue that, on a global scale, many food items are poorly characterized ^14^. As more food-related organisms are sequenced and their nutrient contents are quantified, MEDI inferences will become more and more accurate across diverse human populations. Finally, many of the limitations outlined above also extend to dietary questionnaires.

In conclusion, we have developed a data-driven methodology for inferring dietary and nutritional intake from human stool metagenomic data, called MEDI. MEDI provides a viable alternative to questionnaires for assessing dietary intake and it can be readily applied to the large treasure trove of existing metagenomic shotgun sequencing data for which dietary information is not available. By leveraging a common data type that is regularly collected to investigate the composition of the human gut microbiota, MEDI provides a value-addition to any past, present, or future metagenomic study where dietary intake information would prove useful.

## Methods

### Food genome database and mapping hash construction

CSV files describing the FOODB version 1.0 were downloaded and all food items with a mapping to the NCBI Taxonomy Database were extracted. NCBI Taxonomy IDs were mapped onto canonical ranks using taxonkit (version 0.15.0) and searched for in the NCBI GenBank first, followed by a search of the NCBI Nucleotide Database. Manifest files listing all available assemblies were downloaded from NCBI GenBank and each species ID from in the prior step was searched for in the assembly table. All food-derived NCBI taxonomic IDs without a species-level match were then matched at the genus level. Whenever there were multiple potential matches, they were ranked in preference by submission date, preferring most recent to oldest, and by Refseq quality, preferring “reference genome” over “representative genome”. Finally, within this ordering, additional ties were cleared by the assembly type, preferring complete genomes to chromosomes, followed by contigs. All food-derived taxa without a match to NCBI GenBank were then searched in the full NCBI Nucleotide Database for any partial genomic assembly or genes of at least 10,000 base pairs in length, returning the longest contiguous sequences and up to a maximum of 500 records. Similarly, the NCBI Nucleotide Database searches were first performed on the species rank, followed by search on the genus rank, for all non-identified taxa. All matched assemblies were then downloaded in parallel using the NCBI eFetch API, annotated with the corresponding NCBI taxonomic ID in Kraken annotation format, and compressed. Food information, macronutrient composition, energy content and detailed metabolite profiles were extracted from the FOODB data for each matched taxon. A new Kraken2 database was built using the same initial taxonomy dump as used in the previous ID matching with the *kraken2-build* command. This database was first filled with decoy sequences, comprising all complete genomes from bacteria, archaea, viruses, and the human reference genome (Gchr38), sourced from NCBI GenBank. The raw hash table of food-related genomic sequences and decoy genomes was approximately 350GB in size. The database was then indexed with Bracken for 100 and 150bp reads.

### Metagenomic data download and preprocessing

Raw metagenomic shotgun sequencing data was downloaded from the NIH Sequence Read Archive for all data sets using the SRA download pipeline from https://github.com/gibbons-lab/pipelines. Preprocessing was performed using FASTP ^54^, trimming the first 5 bases at the 5’ end and using a sliding window trimming on the 3’ end with a cutoff of a quality score of 20. Reads shorter than 50bps after trimming were discarded from the analysis. Quality control summaries were inspected using MultiQC ^55^, verifying that samples were free of any remaining sequencing adapters and that the insert size in paired-end data was correct.

### Metagenomic mapping and read counting

The MEDI pipeline was implemented as a set of Nextflow workflows ^56^ and is available at https://github.com/gibbons-lab/medi. For all mapping performed in this publication, Kraken2 was run with memory mapping turned on to parallelize database loading across all running mapping processes. This allowed for instantaneous parallelized reading of the large mapping index and led to much lower amortized computation time for individual processes. Kraken2 was run with a default confidence cutoff of 0.3. In order to improve mapping accuracy, we also combined this with an additional post-mapping filter that worked specifically on the canonical ranks of reads and individual kmers classified within each read (Fig. 2A). Here, reads were filtered based on cutoffs for consistency, mapping entropy, and multiplicity. Consistency denotes the fraction of kmer-level taxonomy assignments that are contained in the final read classification. Hence, let *S_i_* = {*s_r_*}*_i_* be the set of taxonomic identifiers assigned to read i for all ranks r and let *K_i_* = {*k_j_*} be the least common ancestor (LCA) taxonomy assignments for each classified kmer j in read i. Then the consistency of read i is given by *C_i_* = |{*k_j_*∈*S_i_*}|/|*K_i_*|. A consistency of 1 means that all individual kmer assignments fall onto a single taxonomic path, whereas a low consistency means that many individual kmer assignments lie on conflicting branches. Multiplicity is the number of unique kmers assignments at the same rank r as the read assignment, *M_i_* = |{*k_j_*|rank(*k_j_*)=*r*}|. Finally, mapping entropy is the Shannon index of the kmer assignments on the same rank as the final read assignment. Thus, if *p_r_*(*k*) is the relative frequency of taxon k at rank r (relative abundance of kmer classifications), then the mapping entropy is given as 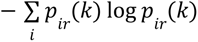. Scoring and filtering methods were implemented in the *architeuthis* software (https://github.com/cdiener/architeuthis) in the Go programming language (https://golang.org). After Kraken2 read-level classification, we removed all reads with consistency smaller 0.95, entropy larger 0.1, and multiplicity larger than 4. This typically removed about 10% of the classified reads from Kraken2. Final abundance estimation was performed with Bracken, using a minimum clade abundance of 10 reads to avoid redistribution to taxa with very low occurrence.

### Infant metagenomic time series

Raw sequencing data was downloaded and processed from the NIH Sequence Read Archive project PRJNA473126 as described above. Preprocessed FASTQ files were then analyzed with the MEDI pipeline, described earlier. Diet summaries from the metadata provided on SRA were used to identify infant-timepoint pairs where a specific food group was consumed. The onset of solid food consumption was the first timepoint for each infant where either Fruits, Meat, Cereal, or Sweets were listed in the diet summary. Associations of total food reads against age were run in a linear mixed effects model using log10(reads + 1 / total reads) as the dependent variable and infant age as a fixed effect with random intercepts and slopes for individual infants. The delivery mode (categorical: vaginal birth vs. cesarean) and abundance of bacterial reads (log10[bacteria reads + 1 / total reads]) were used as covariates.

### Integrative Human Microbiome (iHMP) Project IBD data

A list of individual Run IDs were obtained from the NIH SRA bioproject PRJNA398089, keeping only healthy controls. Metadata for the samples was downloaded from https://ibdmdb.org keeping only those raw FASTQ files with a match to the metadata. Samples were processed as described above and analyzed with the MEDI pipeline. All iHMP raw sequencing files were also processed by an in-house metagenomics pipeline that uses the Kraken2 and Bracken with the default database (not containing foods)e (https://github.com/gibbons-lab/pipelines). BRACKEN estimated species abundances for were averaged across all samples and only taxa with at least 100 reads on average were kept. This yielded an average abundance profile for non-food associated taxa in the human gut.

### METACARDIS data

Raw sequencing data was downloaded and processed from the NIH Sequence Read Archive project PRJEB37249 as described above. Metadata were obtained from Supplemental Table 14 from Fromentin et al. (2022) ^43^.

### Simulated ground truth data

Reference genomes for all bacteria and archaea in the decoy data used in the Kraken2 mapping database were downloaded along with a 1000 genome project human genome assembly ^57^. This human genome assembly was used instead of the one in the Kraken2 database to test for the effect of having human sequence fragments that may not have been represented in the decoy database. A background relative abundance distribution for bacterial, archaeal, and viral taxa was established using the iHMP healthy cohort, as described above. The respective NCBI taxonomy IDs for each species in the average abundance profile were used to match organisms to reference genomes from the NCBI RefSeq database and download the assemblies in FASTA format. Reads of 150bps length were sampled using DWGSIM (https://github.com/nh13/DWGSIM) using a uniformly decreasing (5’ to 3’) error rate of 0.001-0.005 for the forward reads and 0.05-0.01 for the reverse reads to a final depth of 10 million paired-end reads per sample. For this, reads were sampled from each individual reference genome to a final n of r_i_*10,000,000 where r_i_ is the relative abundance of taxon i in the background abundance profile. A negative sample was generated by sampling from the background. Positive samples were generated by repeatedly choosing 10 random food species and adding 1 million reads to 9 million reads of the background (10% final food read abundance) using the food genomic sequences downloaded during database construction and read sampling as described above. Individual food abundances were staggered by a natural log level for each of the 10 food items, yielding 10 log levels of abundance variation, ranging from less than 10 to more than half a million reads across the 10 food species. The simulation of food-positive samples was repeated 16 times (16 random sets of 10 foods). After sampling, reads were shuffled and quantified using the MEDI mapping pipeline described before. The false positive mapping rate was quantified as the fraction of reads that were assigned to a food item not present in the sample. Mapping accuracy was quantified as the Pearson product-moment correlation r of expected relative read abundance versus the observed read abundance.

## Acknowledgements

Research reported in this publication was supported by the National Institute of Diabetes and Digestive and Kidney Diseases (NIDDK) of the National Institutes of Health (NIH) under award number R01DK133468 (to SMG) and by a Global Grants for Gut Health Award from Nature Portfolio and Yakult (to SMG). CD was supported by the Austrian FWF Cluster of Excellence: Microbiomes Drive Planetary Health.

---

**Figure S1.**
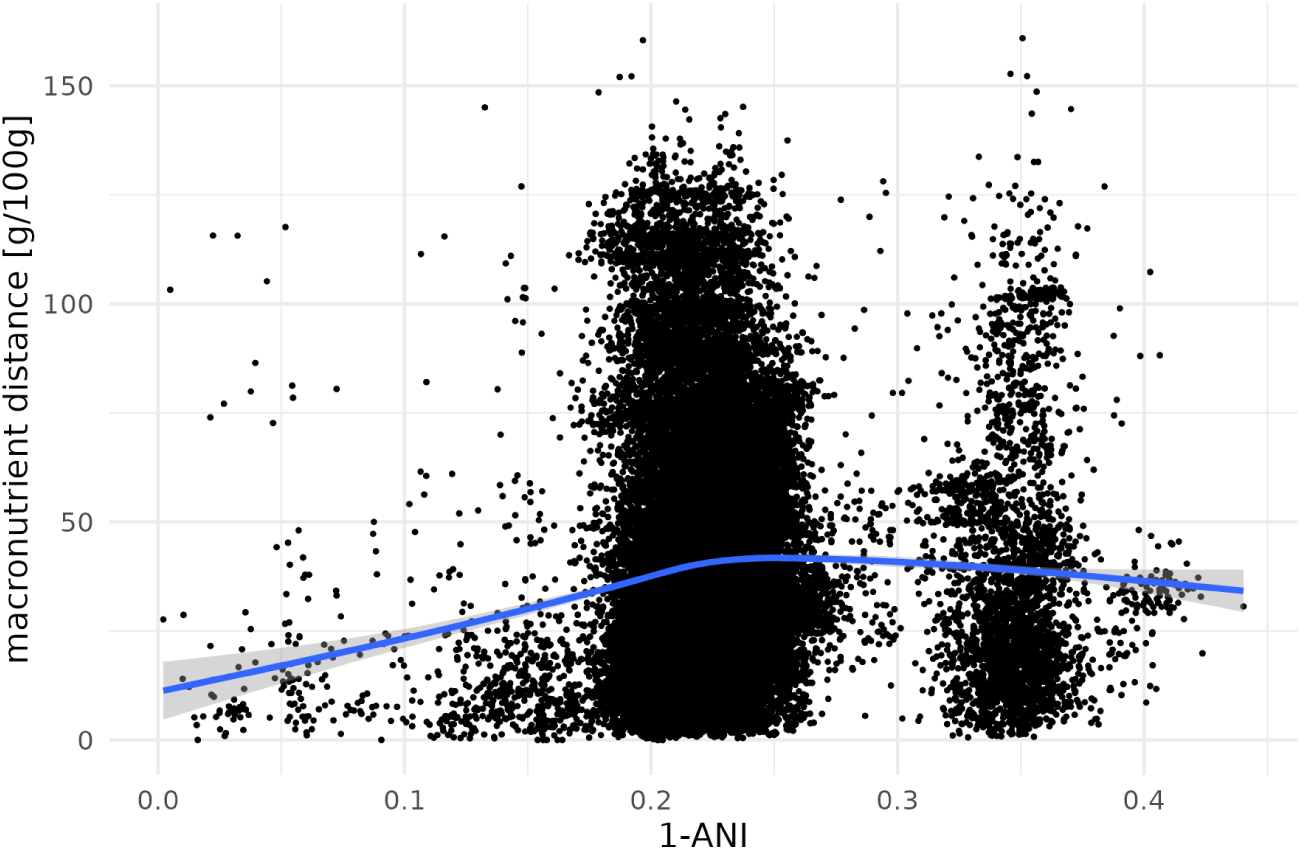
Genomic distance (1 - ANI) vs. macronutrient distance (euclidean). The blue line denotes a smooth spline regression and shaded area denotes the 95% confidence interval of the spline regerssion.

**Figure S2.**
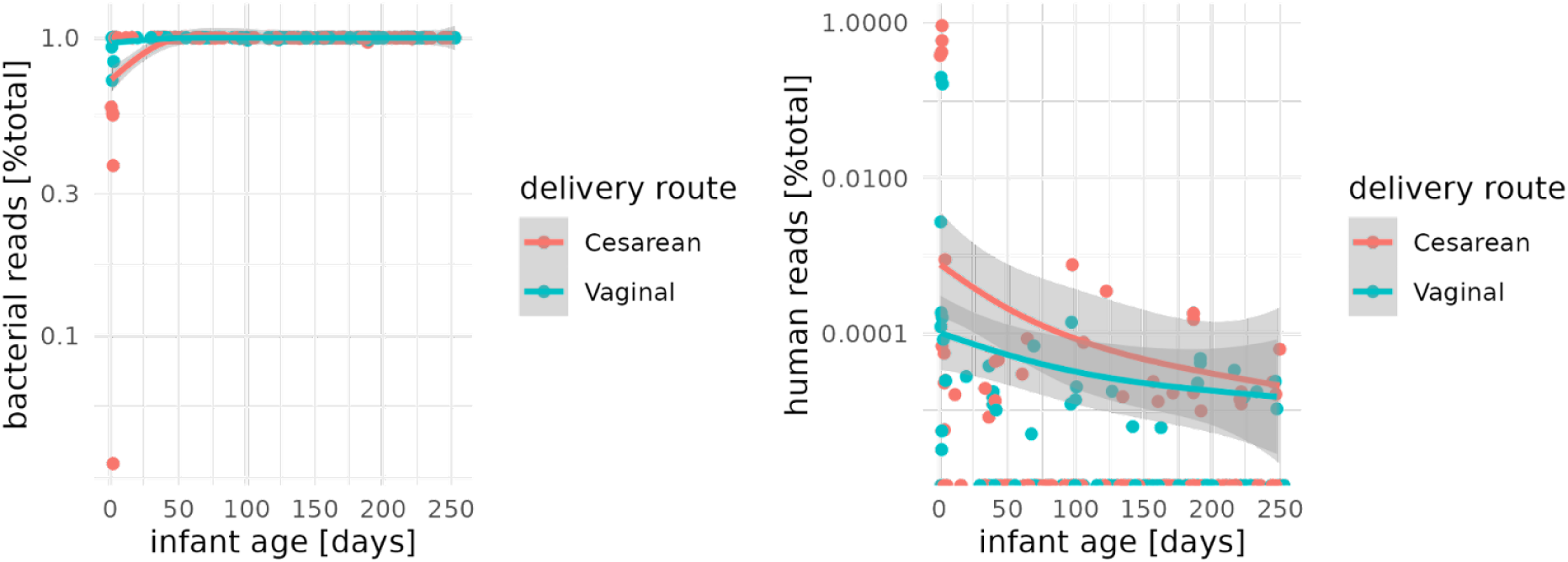
Relative abundance of bacterial and human reads across infant timeseries, colored by delivery route.

**Figure S3.**
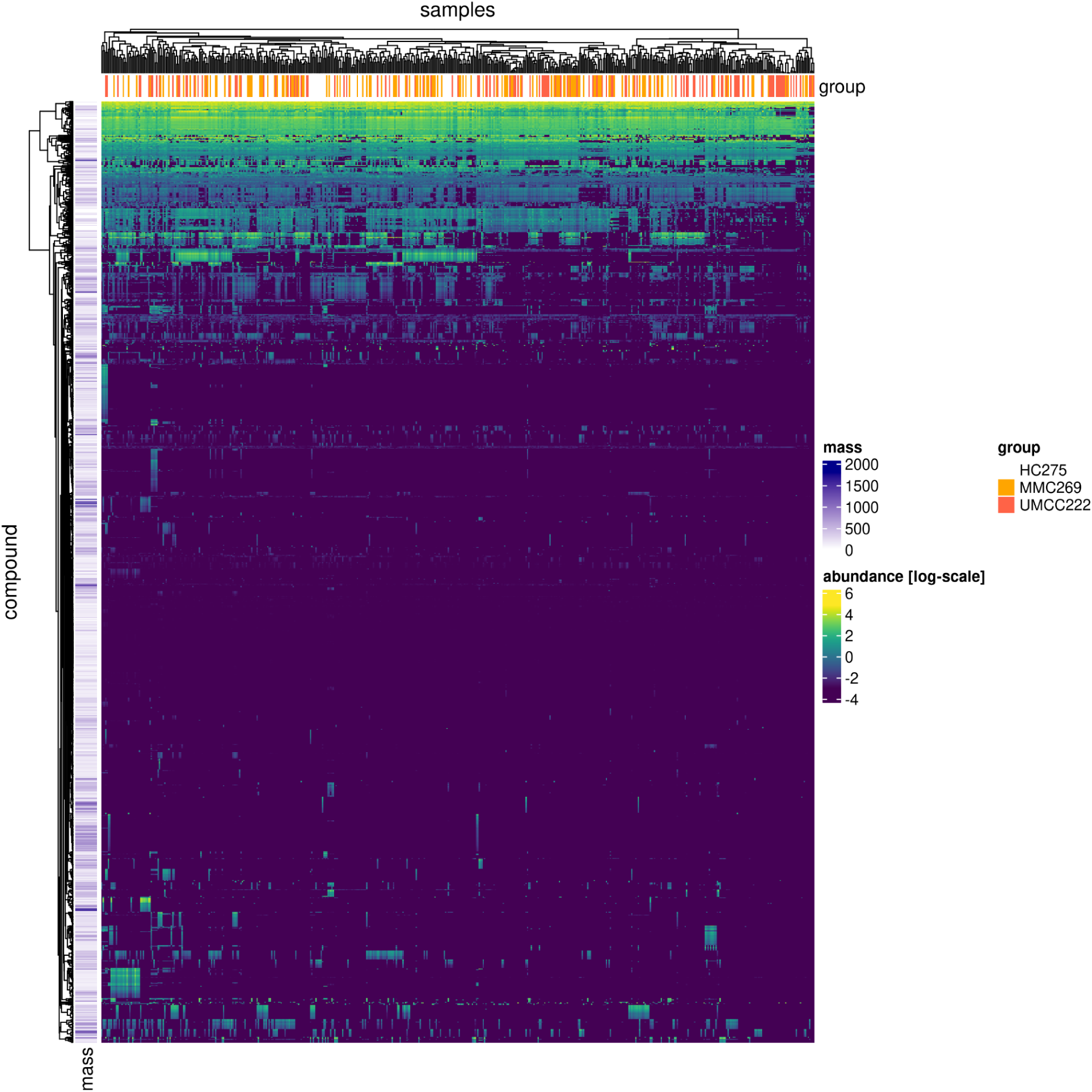
MEDI dietary intake estimates were associated with metabolic health. (A) Abundances per 100 g portion for 1703 compounds across a cohort of 533 metabolically healthy and unhealthy individuals from the METACARDIS cohort. Fill colors denote abundance per standard portion (mg/100g). Column annotations denote metabolic health status from the original METACARDIS cohort (HC - healthy cohort, MMC - IHD metabolically matched cohort, UMMC - untreated metabolically matched cohort). Here, MMC and UMMC denote disease-free but metabolically unhealthy groups. Row annotations denote the monomer mass of the compound (in g/mol).

